# Positioning Genomic Features in Biomedical Knowledge Graphs using the Homo sapiens Chromosomal Location Ontology for GRCh38 (HSCLO38)

**DOI:** 10.1101/2024.02.15.580505

**Authors:** Taha Mohseni Ahooyi, Benjamin Stear, Deanne M. Taylor

## Abstract

The Homo sapiens Chromosomal Location Ontology for GRCh38 (HSCLO38) represents a knowledge-graph-ready framework for connecting genomic features at multiple resolutions. We present the methodology behind the development of HSCLO38 and its integration with current genomic standards for application in biomedical research. We explore the performance and scalability of HSCLO38 in specific use cases in handling large-scale genomic data in a biomedical knowledge graph.

## Introduction

Knowledge graphs (KGs) are emerging as a useful way to integrate and analyze heterogeneous biomedical data. Genomic data could be incorporated and integrated using KGs, but one consideration is supporting differing experimental resolutions from entire chromosomes to individual base pairs. Integrating and analyzing genomic features through basic numerical coordinates in large-scale biomedical KGs can lead to significant computational demands. We were interested in developing a system to reduce computational overhead but still be able to analyze genomic features in any biomedical knowledge graph by integrating data across different experimental resolution levels. To answer this challenge, we created the Homo sapiens Chromosomal Location Ontology for GRCh38 (HSCLO38). HSCLO38 is represented as an ontologized genomic coordinate binning schema. HSCLO38’s purpose is to simplify the integration of genomic experimental data at different resolution scales within any biomedical knowledge graph. Other knowledge graphs have been designed for identifying, mapping, or analyzing genomic features (Feng et al. 2022), however, HSCLO38 is a system designed to be utilized by any knowledge graph interested in utilizing a method for rapid integration of genomic features by GRCh38.

We outline the development of HSCLO38, detail its integration with established genomic standards such as GENCODE, and discuss its application in connecting genomic features. We have used this ontology to incorporate genomic datasets of different resolutions within KGs, including Hi-C physical contact regions, ATAC-seq chromatin accessibility data, functional DNA elements like genes and regulatory regions, and base-pair level features such as single nucleotide variants within regulatory elements.

## Results

HSCLO38 defines chromosomal locations within the GRCh38 release for chr 1-22, X, Y, and M (**Figure 1**). It provides hierarchical relationships across five genomic resolution levels: whole chromosome, 1 megabase pair (Mbp), 100 kilobase pairs (kbp), 10 kbp, and 1 kbp. Each node within these class levels is interconnected to its scale parent and the immediate neighbors on either side to support mapping and association between genomic datasets and features. For example, the 1kbp element HSCLO38:chr1.20517001-20518000 is_a connected to its “scale parent” HSCLO38:chr1.20510001-20520000 as well as to the 5’ neighbor HSCLO38:chr1.20516001-20517000 and 3’ neighbor HSCLO38:chr1.20518001-20519000. Using human genome version GRCh38, the HSCLO38 schema results in 3,431,155 nodes and 6,862,195 relationships (**Table 1**).

**Table 1:**
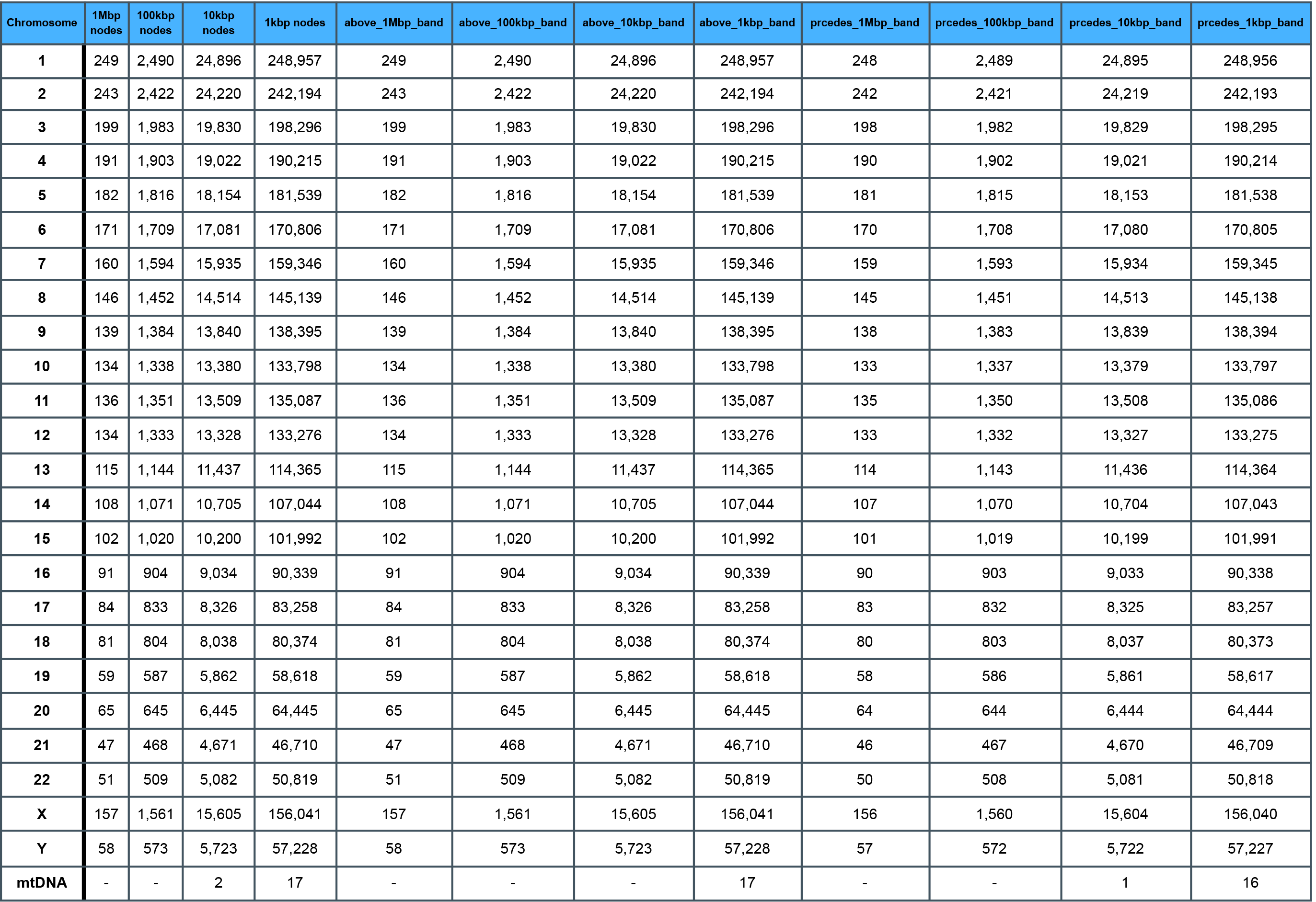
HSCLO38 node and relationship statistics.

**Figure 1:**
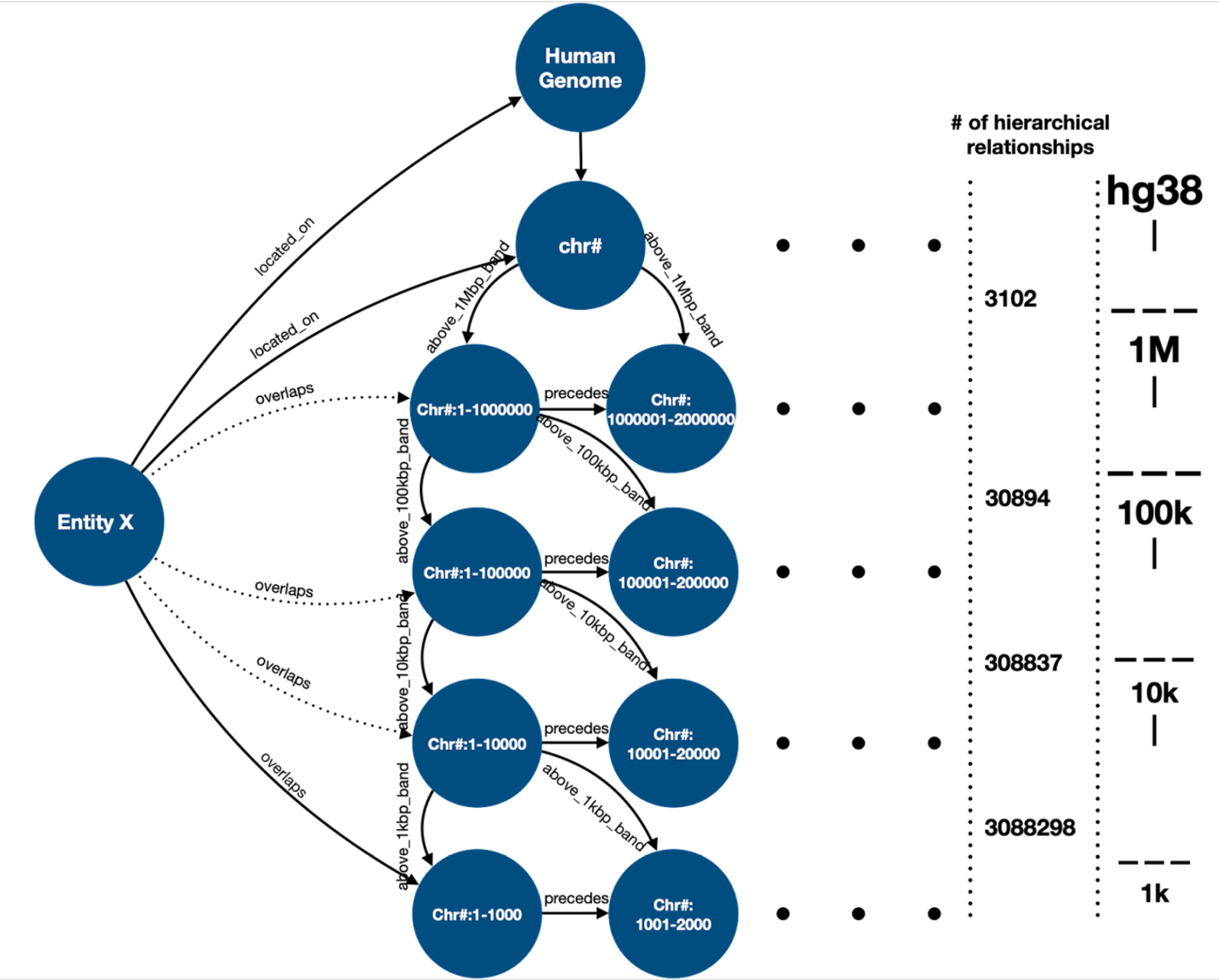
The schema of the Chromosome Region ontology developed for Petagraph. Entity X could be any chromosomal feature, including chromosomal bands, genes, exons, introns, regulatory elements, QTLs, variants, accessible chromatin regions, viral integration sites, human endogenous retroviruses, transposons, tandem repeats, chromosomal contact regions, TADs, telomere, centromeres, and any other type.

We provide a use case for linking biodata at different resolutions to demonstrate the practical application of HSCLO38 in knowledge organization and discovery. A researcher may be interested in identifying genes within large-scale chromatin organization features, such as Hi-C data hosted by the 4DN project (Dekker et al. 2023). We began by importing HSCLO38 into a biomedical KG (Stear et al. 2023), and then creating edges in the KG to link all gene nodes from GENCODE v41 (Harrow et al. 2012) to their respective 1kbp HSCLO38 nodes. We then created edges for the chromosomal loops from a set of files at the 4DN project (Dekker et al. 2023) to their respective 1kbp locations in HSCLO. Using a Cypher query in the Neo4j v5 environment, we retrieved the overlap in 1kbp nodes between the spans of the GENCODE gene definitions and the start and end points of the 4DN loops.

**Figure 2** shows the distribution of 4DN loop sizes (**Figure 1A**) and the number of GENCODE-defined genes overlapping the 4DN dataset loops (**Figure 1B & 1C**). The mode of the distribution occurs at 2 genes. Further analysis of this data reveals ∼3000 loops (from 21 4DN dotcall files) that overlap at least 10 genes per 100kbp of the loop length.

**Figure 2:**
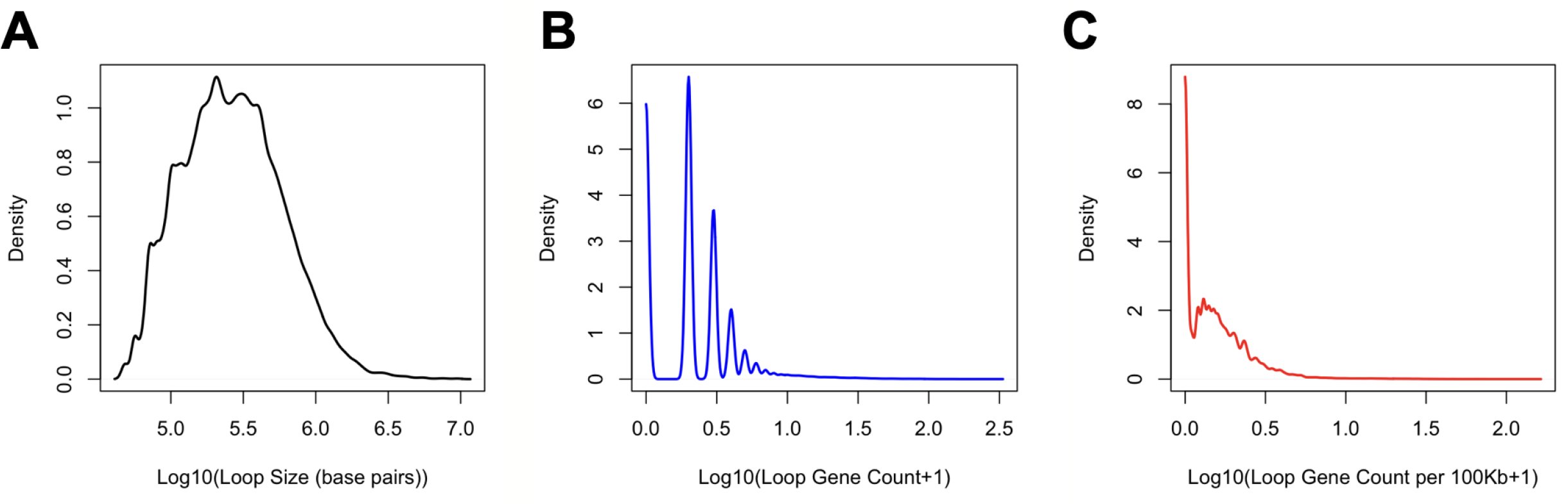
Loop size (A) and gene count distribution (B and C) derived from the intersection of 4DN loop and GENCODE genes entities identified through their connection to HSCLO38

To explore the biological relevance of this analysis, we performed functional annotation of the gene list from the loop with the highest number of overlapping genes (4DNFI3GNGT17.chr2.150000-170000.chr2.250000-270000). The analysis provides the top 10 enriched pathways (**Table 2**), top 10 DisGenNet diseases (**Table 3**), and top 10 MSigDB cell types (**Table 4**), implying the disruption in the loop structure and subsequently the expression regulation of the overlapping genes could be associated with developmental disorders primarily related to muscular development.

**Table 2:** Top 10 pathways associated with genes overlapping loop 4DNFI3GNGT17.chr2.150000-170000.chr2.250000-270000.

| Term | Category | Description | Count | % | Log10(P) | Log10(q) |
| --- | --- | --- | --- | --- | --- | --- |
| WP5224 | WikiPathways | 2q37 copy number variation syndrome | 16 | 8.99 | -15.39 | -11.04 |
| GO:0007517 | GO Biological Processes | muscle organ development | 14 | 7.87 | -8.27 | -4.35 |
| GO:0007423 | GO Biological Processes | sensory organ development | 18 | 10.11 | -7.96 | -4.35 |
| GO:0048598 | GO Biological Processes | embryonic morphogenesis | 18 | 10.11 | -7.89 | -4.35 |
| M160 | Canonical Pathways | PID AVB3 INTEGRIN PATHWAY | 8 | 4.49 | -7.88 | -4.35 |
| GO:0003012 | GO Biological Processes | muscle system process | 13 | 7.30 | -7.64 | -4.25 |
| GO:0007610 | GO Biological Processes | behavior | 17 | 9.55 | -6.81 | -3.68 |
| hsa05205 | KEGG Pathway | Proteoglycans in cancer | 10 | 5.62 | -6.39 | -3.32 |
| GO:0010035 | GO Biological Processes | response to inorganic substance | 15 | 8.43 | -6.25 | -3.21 |
| GO:0009725 | GO Biological Processes | response to hormone | 18 | 10.11 | -6.05 | -3.06 |

**Table 3:** Top 10 DisGenNet abnormalities associated with genes overlapping loop 4DNFI3GNGT17.chr2.150000-170000.chr2.250000-270000.

| Term | Description | Count | % | Log10(P) | Log10(q) |
| --- | --- | --- | --- | --- | --- |
| C0035229 | Respiratory Insufficiency | 23 | 13.00 | -18.00 | -13.00 |
| C0013421 | Dystonia | 24 | 13.00 | -15.00 | -11.00 |
| C0454644 | Delayed speech and language development | 26 | 15.00 | -15.00 | -11.00 |
| C0541794 | Skeletal muscle atrophy | 20 | 11.00 | -15.00 | -11.00 |
| C1145670 | Respiratory Failure | 20 | 11.00 | -14.00 | -11.00 |
| C4552811 | Generalized Muscle Weakness, CTCAE | 14 | 7.90 | -14.00 | -10.00 |
| C1854301 | Motor delay | 21 | 12.00 | -14.00 | -10.00 |
| C0005745 | Blepharoptosis | 25 | 14.00 | -14.00 | -10.00 |
| C0011168 | Deglutition Disorders | 21 | 12.00 | -14.00 | -10.00 |
| C0033377 | Ptosis | 25 | 14.00 | -14.00 | -10.00 |

**Table 3:** Top 10 MSigDB cell types associated with genes overlapping loop 4DNFI3GNGT17.chr2.150000-170000.chr2.250000-270000.

| Term | Description | Count | % | Log10(P) | Log10(q) |
| --- | --- | --- | --- | --- | --- |
| M40176 | DESCARTES FETAL EYE SKELETAL MUSCLE CELLS | 9 | 5.10 | -6.20 | -3.80 |
| M40093 | DESCARTES MAIN FETAL SKELETAL MUSCLE CELLS | 9 | 5.10 | -5.70 | -3.50 |
| M39233 | LAKE ADULT KIDNEY C14 DISTAL CONVOLUTED TUBULE | 8 | 4.50 | -4.50 | -2.50 |
| M39309 | CUI DEVELOPING HEART COMPACT VENTRICULAR<br>CARDIOMYOCYTE | 5 | 2.80 | -4.40 | -2.50 |
| M39093 | ZHONG PFC C6 DLX5 GAD1 GAD2 POS INTERNEURON | 3 | 1.70 | -4.10 | -2.20 |
| M39253 | MENON FETAL KIDNEY 3 STROMAL CELLS | 5 | 2.80 | -3.90 | -2.10 |
| M39209 | HAY BONE MARROW STROMAL | 14 | 7.90 | -3.70 | -2.00 |
| M40178 | DESCARTES FETAL EYE LENS FIBRE CELLS | 5 | 2.80 | -3.70 | -2.00 |
| M40180 | DESCARTES FETAL EYE STROMAL CELLS | 5 | 2.80 | -3.70 | -2.00 |
| M39050 | MANNO MIDBRAIN NEUROTYPES HPERIC | 14 | 7.90 | -3.60 | -1.90 |

## Discussion

The implementation of HSCLO38 offers a knowledge-graph-friendly approach to integrating heterogeneous, multi-resolution biomedical data, providing enhanced accessibility to genomic information. Through an example use case, we showed that HSCLO38 can provide an easy-to-use bridge between different experimental resolutions such as DNA loops and genes, and therefore can facilitate the process of knowledge extraction within relational databases such as graph knowledge graphs.

## Methods

R code was written to parse the GRCh38 coordinate files into binned locations (nodes) connected as an ontology by size scale. HSCLO nodes are defined at 5 resolution levels; chromosomes, 1 Mbp, 100 kbp, 10 kbp, and 1kbp with each level connecting to the lower level with edge names “above_(resolution level)_band” (e.g. “above_1Mbp_band”, “above 1_kbp_band”) and nodes at the same resolution level are connected through edge names “precedes_(resolution level)_band” (e.g. “precedes_10kbp_band”).

Analysis for density estimation was performed in R (RStudio 2022.12.0.353 and R v4.2.2)

Functional annotation of the gene list associated with the chromosomal loop with the highest number of genes per 100kbp loop length was done using Metascape (Zhou et al. 2019).

## Data Availability

A knowledge-graph-ready edgelist (triple format) can be found on the HSCLO38 project page at the OSF website: https://osf.io/pe8v7/.

## Code Availability

The code used to generate and query HSCLO38 is available in a public repository: https://github.com/TaylorResearchLab/HSCLO/tree/main/HSCLO38

## Acknowledgments

We would like to acknowledge useful feedback and discussion on HSCLO38 implementation and support within the Unified Biomedical Knowledge Graph (UBKG) from J. Alan Simmons and Jonathan C. Silverstein at the Department of Biomedical Informatics at the School of Medicine, The University of Pittsburgh. Work on HSCLO38 was partially supported by the NIH Common Fund Data Ecosystem Partnership award to the Kids First Data Resource Center.

## Author Contributions

TAM wrote the code, and provided analyses and figures. TAM, DMT and BJS wrote and edited the paper. TAM, DMT, and BJS designed the HSCLO38 schema. TAM and BJS implemented HSCLO38 in the knowledge graphs. DMT conceived of, guided, and funded work on HSCLO38. JCS and JAS provided

## Version

15 Feb 2022 a

